# Schistosomiasis Induces Persistent DNA Methylation and Tuberculosis-specific Immune Changes

**DOI:** 10.1101/255125

**Authors:** Andrew R. DiNardo, Tomoki Nishiguchi, Emily M. Mace, Kimal Rajapakshe, Godwin Mtetwa, Alexander Kay, Gugu Maphalala, W. Evan Secor, Rojelio Mejia, Jordan S. Orange, Cristian Coarfa, Kapil N. Bhalla, Edward A Graviss, Anna M. Mandalakas, George Makedonas

**Author notes:** Co-last authors contributing equally (Anna M. Mandalakas, George Makedonas). **Corresponding author:** Andrew DiNardo, MD, Feigin Center, suite 630, 1102 Bates, Houston, Texas, 77030.

## Abstract

Epigenetic mechanisms, like DNA methylation, determine immune cell phenotype. To understand the epigenetic alterations induced by helminth co-infections, we evaluated the longitudinal effect of ascariasis and schistosomiasis infection on CD4+ T cell DNA methylation and the downstream tuberculosis (TB)-specific and BCG-induced immune phenotype. All experiments were performed on human primary immune cells from a longitudinal cohort of recently TB-exposed children. Compared to age-matched uninfected controls, children with active *Schistosoma haematobium* and *Ascaris lumbricoides* infection had 751 differentially DNA methylated genes with 72% hyper-methylated. Gene ontology pathway analysis identified inhibition of IFN-γ signaling, cellular proliferation, and the Th1 pathway. Targeted RT-PCR after methyl-specific endonuclease digestion confirmed DNA hyper-methylation of the transcription factors *BATF3, ID2, STAT5A, IRF5, PPARg, RUNX2, IRF4* and *NFATC1* and cytokines or cytokine receptors *IFNGR1, TNFS11, RELT* (TNF receptor), *IL12RB2* and *IL12B* (p< 0.001; Sidak-Bonferroni). Functional blockage of the IFN-γ signaling pathway was confirmed with helminth-infected individuals having decreased up-regulation of IFN-γ-inducible genes (Mann-Whitney p < 0.05). Hypo-methylation of the IL-4 pathway and DNA hyper-methylation of the Th1 pathway was confirmed by antigen-specific multidimensional flow cytometry demonstrating decreased TB-specific IFN-γ and TNF and increased IL-4 production by CD4+ T cells (Wilcoxon signed rank P <0.05). In S.haematobium infected individuals, these DNA methylation and immune phenotypic changes persisted at least six months after successful deworming. This work demonstrates that helminth infection induces DNA methylation and immune perturbations that inhibit TB-specific immune control and that the duration of these changes are helminth-specific.

## Introduction

The immune system has multiple mechanisms to remain responsive to changing environments. It is now clear that both the innate and adaptive immune response use epigenetic mechanisms of gene regulation to maintain long-term phenotypes. For example, monocytes “trained” with β-glucan or BCG undergo epigenetic chromatin alterations that alter TNF and IL-6 production and improve *Staphylococcus*, *Candida* and mycobacterial killing(1, 2). Long-term phenotypic changes in adaptive immunity are similarly epigenetically mediated. The RAG-mediated T cell receptor (TCR) rearrangement requires hypo-methylation of *RAG* and other critical genes(3). After TCR rearrangement, T cells remain responsive to changing environments via epigenetic modifications that polarize CD4^+^ T cells into phenotypic subsets (Th1, Th2, Th9, Th17, Tregs, Th22, and Tfh)(4-6). To differentiate into a Th1 CD4^+^ T cell, the *IFNG* promoter is de-methylated and the *IL4* locus undergoes repressive histone modifications(7-9). While previously thought to be immutable, epigenetic mechanisms have recently been described that allow T cells to shift between Th1 and Th2 phenotypes depending on environmental stimuli(10). Understanding the duration and mechanisms of epigenetic mediated immune polarization is critical to predicting and reversing detrimental immune perturbations.

Helminths are a heterogeneous group of parasites estimated to affect a quarter of the world’s population(11, 12). While most infections are subclinical, they can induce growth retardation, nutritional deficiencies and profoundly perturb immunogenicity(12-14). The helminth-induced immune response is characterized by tolerance and a Type 2 immune polarization(15, 16) with decreased IFN-γ, IL-12 and IL-17(17), and increased T regulatory cells (Tregs)(18), IL-4, IL-5, IL-10(19) and TGF-*β(20)*. This profound Type 2 and immune-tolerant phenotype undermines the Type 1 immune response to cholera, salmonella, tetanus, and BCG(21-25) and is the suspected rational for increased TB disease progression among helminth infected individuals(11, 26-28). By contrast to the Type 2 immune response required to expel helminths, *Mycobacterium tuberculosis (Mtb*) infection requires a Type 1 polarized CD4-macrophage immune axis for effective host response.

Individuals with IL-12-IFN-γ axis defects due to Mendelian Susceptibility to Mycobacterial Diseases (MSMD), HIV and TNF-inhibitors are prone to disseminated BCG, TB disease progression and infection with other intracellular pathogens(29, 30). Stimulation of macrophages with TNF and IFN-γ induces intracellular killing via phago-lysosome maturation and nitric oxide (NO) production, while IL-4 induces a diametrically opposed reaction to direct macrophages away from NO and towards collagen synthesis(31). Therefore, when macrophages are polarized with IL-4 they exhibit decreased phago-lysosome maturation, diminished NO production and reduced *Mtb* killing(32-35).

In the mêlée of host-pathogen interactions, pathogens induce epigenetic changes to subvert host immunity. HIV induces DNA methyltransferases (DNMT) expression that hyper-methylates *IFNG* and *IL2*. HIV also hypo-methylates *PD1*, resulting in decreased IFN-γ and IL-2 while increasing PD-1*(36-39)*. Similar to other intracellular pathogens(40-42), *Mtb* induces epigenetic changes that benefit mycobacterial survival by altering macrophage (MΦ) antigen-presentation and inflammasome function(43-48). CD4^+^ T cell cytokine production is critical for macrophage polarization and *Mtb* control(49, 50). In murine studies, schistosomiasis induces epigenetic changes that drive alternative activated macrophages(7) and a Th2 CD4^+^ T cell phenotype characterized by DNA hyper-methylation of the *IFNG* promoter(8). Therefore, to elucidate the longitudinal epigenetic-immune effects of helminth co-infection, we evaluated the effect of *Schistosoma haematobium* and *Ascaris lumbricoides* on CD4^+^ T cell DNA methylation and TB-specific immune phenotype. These studies indicate that *Schistosoma* and *Ascaris* induce global changes in host DNA methylation and immune phenotypic changes in CD4^+^ T cells. For *S. haematobium*, these host DNA methylation and immune phenotypic perturbations are relatively stable and persist at least six months after successful anti-helminthic therapy.

## Patients and Methods

### Cohort

Asymptomatic children 6 months to 15 years of age with recent intense TB exposure (as quantified by a standardized TB exposure score; Supplemental Table I) were enrolled after obtaining assent and written informed consent of their parent or caregiver. As per national guidelines, participants were empirically dewormed with albendazole and treated with isoniazid preventative therapy (IPT) if <5 years of age or HIV-infected. Individuals with HIV infection were excluded from this analysis. All individuals were BCG-vaccinated as confirmed by BCG scar and/ or vaccine card. Participants completed clinical evaluation at 2 and 6 months for TB symptoms and provided stool and urine for helminth evaluation. Blood was collected for helminth serology and peripheral blood mononuclear cell (PBMC) collection at each visit. Stool and urine were collected for parasite microscopy and PCR at each visit. The protocol was reviewed and approved by the Baylor College of Medicine Children’s Foundation-Swaziland (BCMCF-SD), Baylor College of Medicine and the Swaziland Scientific and Ethics Committee. The protocol was also reviewed at the Centers for Disease Control and Prevention, which issued a non-engaged determination. All methods were performed in compliance with institutional and international guidelines for the Protection of Human Subjects in concordance with the Declaration of Helsinki.

### Helminth Status

To comprehensively describe a child’s helminth status, serology, ova and parasite microscopy (O&P) and helminth-specific multi-parallel quantitative real-time PCR were performed. Schistosomiasis serology was performed by standard ELISA. In brief, 96-well ELISA plates were coated with 2µg/ml *Schistosoma mansoni* soluble egg antigen (SEA) in 0.1M sodium bicarbonate buffer (pH 9.6) for 2 hours followed by washing with PBS + 0.3% Tween 20, blocking with PBS/Tween containing 5% dry milk powder. Plates were again washed and incubated with 1:50 dilution of participant plasma in PBS/Tween/milk for 30 minutes. Plates were again washed and incubated with a 1:50,000 dilution of horseradish peroxidase conjugated mouse anti-human IgG (Southern Biotech, Birminham, AL) for 30 minutes. After a final wash, SureBlue^TM^ TMB (Kirkegaard & Perry, Gaithersbug, MD) substrate was added to plates for 5 minutes and the reaction was stopped using 1N H_2_SO_4_. Plates were read at 450nm on a Molecular Devices (Sunnyvale, CA) microplate reader and patient results were compared to a standard curve consisting of sera from confirmed schistosomiasis patients that was included on each plate using SOFTmaxPro software (Molecular Devices).

Stool microscopy was performed using the McMaster protocol(51). Two grams of stool were added to 28mL of NaCl and sugar flotation solution and homogenized by aggressive hand mixing followed by straining and viewing on a McMaster chamber slide. Urine microscopy was performed after 400g centrifugation.

Multi-parallel helminth qPCR was performed as previously described(52). DNA was extracted from 50mg of stool using the MP FastDNA for Soil Kit. For urine DNA extraction, the concentrated pellet was heated at 90^o^C for 10 minutes followed by DNA isolation using the MP FastDNA kit. From stool, helminth-specific PCR was performed for A. lumbricoides, Necator americanus, Ancylostoma duodenale, Trichuris trichiura and Strongyloides stercoralis using primers and FAM-labeled minor groove binding probes (Supplemental table II). Similarly, qPCR specific for *S. haematobium* was performed from the DNA isolated from urine with organism specific primers and probes (Supplemental Table II). The qPCR reactions were performed in duplicate with 2 µL of stool or urine derived template DNA, 900 nM primers, 250 nM probe and 3.5 µL of TaqMan Fast MasterMix and acquired on a QuantStudio3 in Mbabane, Swaziland after 40 cycles. Quantification was performed using a standard curve consisting of 10-fold serial dilutions of purified plasmid DNA containing the target sequences.

Composite helminth status was defined according to serology, microscopy and PCR status. Active helminth infection was defined as microscopy and/ or PCR positive. Remote infection was classified as serology positive, but microscopy and PCR negative both at baseline and two and six-month follow-up. Uninfected controls were microscopy, PCR and serology negative.

### DNA Methylation

To evaluate CD4^+^ T cell DNA methylation, CD14 positive selection was followed by CD4 negative selection (StemCell) to obtain >93% CD4^+^ T cell population determined by flow cytometry. Cellular DNA was then isolated using the DNeasy Lymphocyte protocol (Qiagen). To perform the DNA MethylEPIC array, Genomic DNA (gDNA) from CD4^+^ T cells was quantified using a Qubit 3.0 fluorometer (Thermofisher Scientific, MA, USA). gDNA quality was analyzed using Agilent 4200 Tape Station System (Agilent Technologies, Inc. CA, USA) for integrity check and Nanodrop ND1000 spectrophotometer for purity check. gDNA (250–750 ng) was treated with sodium bisulphite using the Zymo EZ DNA Methylation Kit (Zymo Research Corp., CA, USA)). We followed the Illumina recommended incubation conditions during bisulfite conversion to maximize DNA conversion rate. Illumina Infinium HumanMethylation850 (HM850) and HumanMethylationEPIC (EPIC) BeadChip (Illumina, CA, USA) were run on an Illumina iScan System (Illumina, CA, USA) using the manufacturer’s standard protocol.

Validation of the Infinium DNA Methylation EPIC data was performed using real-time PCR after isolated DNA underwent methylation-sensitive or dependent digestion with a DNA MethylationEpitect II kit. The relative percentage of methylated and unmethylated DNA was determined relative to mock and double digested DNA as previously described(53).

### Multi-dimensional Immune Profiling

To identify TB-specific and BCG induced immunity, PBMCs were stimulated with 2.5 µg each of ESAT-6 and CFP-10 overlapping peptide pools or 5µg of BCG sonicate, respectively. BCG and TB-specific immune responses were compared to vehicular control (dimethyl sulfoxide) stimulation and staphylococcal enterotoxin B was used as a positive control. Cell stimulations occurred in the presence of brefeldin A (1µg/mL), monensin (0.7 µg/mL) and CD28/CD49 (2.5 µg/mL). Following overnight (16 hour) stimulation, cells were stained with an amine reactive viability dye (Ghost dye), surface antibodies (CD4, CD8, CD56 and PD-1) and intracellular antibodies (CD3, IFN-γ, TNF, IL-4, IL-10, IL-13, perforin, T-bet, GATA-3, and Ki-67) and acquired on a BD LSR II Fortessa (Supplemental Table II). A minimum of 70% viability was the threshold for credible data. All reported TB-specific or BCG-induced values were subtracted from the background non-stimulated immune response.

### Gene Expression

PBMCs were stimulated overnight with IFN-γ (50ng/mL) or media control and 16 hours later CD4+ T cells were isolated via magnetic bead negative selection as described above. From 100ng of RNA, the Human Immunology v2 probe set was performed using the NanoString nCounter. Background noise was normalized using 8 negative control probes and the geometric mean of positive controls and 15 housekeeping genes.

### Monocyte-Derived Macrophage (MDMΦ) Immune Polarization

Adherent monocytes from healthy control PBMCs were cultured for 7 days at 37^o^C with 10% human AB serum and 10ng/mL M-CSF and GM-CSF. After 7-day conversion to MDMΦ, the cells were polarized with IL-4 20ng/mL or IFN-γ 50 ng/mL for 48 hours and then brought to the BSL-3. The MDMΦs from three healthy controls were incubated with live H37Rv *Mtb* at an MOI of 3:1 overnight, non-phagocytosed extra-cellular bacilli were removed by washing, MDMΦ were lysed with ice cold deonized water, and mycobacteria were cultured on 7H11 plates at 1:5 and 1:25 dilutions. Twenty-one days later, colony-forming units (CFU) were manually counted.

### Statistical Analysis

Flow Cytometry data analysis was performed in FlowJo (TreeStar, Ashland, OR); GraphPad 6.0 (GraphPad Software), and SPICE (available at http://exon.niaid.nih.gov/spice). The non-parametric Mann-Whitney test was used to compare data sets. For multiple comparisons, the Sidak-Bonferroni test was performed. Statistically significant differentially methylated genes were visualized using Ingenuity Pathway Analysis (Qiagen Bioinformatics). IDAT files from methylation data were imported into R statistical system using the Bioconductor (http://www.bioconductor.org) minfi package(54).

Preprocessed and normalized data from minfi was used by Bioconductor limma package to identify the significantly differentiated probes with p-value <0.05 and log2 fold change > 2 or <0.5(54). Statistically significant differentially methylated genes were visualized using Ingenuity Pathway Analysis (Qiagen Bioinformatics). Further gene enrichment analysis was performed by employing the gene set collections compiled by the Molecular Signature Database (MSigDB, http://www.broadinstitute.org/gsea/msigdb/) using the hypergeometric distribution (statistical significance achieved at P < 0.05)(55).

## Results

### Participant demographics

Among the children enrolled who had exposure to microbiologically confirmed pulmonary TB case in the past 6 months, fifty-nine percent of children were female and the median age was 6.1 (IQR 2.7-10.5). The median and mean TB exposure score was 7 (Table I).

**Table 1:**
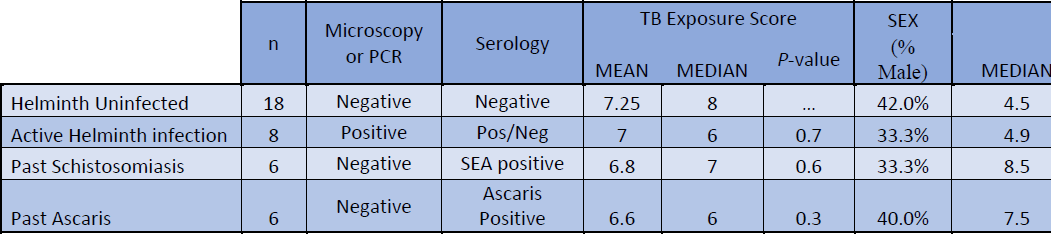
**Demographics** and TB exposure of clinical cohort

### Helminth Infection induces CD4 DNA Methylation Changes that Persist after Schistosomiasis

We evaluated how helminth status altered CD4^+^ T cell DNA methylation. Individuals with current ascaris or schistosome infection (n = 8), confirmed by ova and parasite (O&P) microscopy or by multi-parallel PCR, have unique CD4^+^ T cell DNA methylation signatures compared to uninfected, TB-exposed children (n = 18). Using the DNA Methylation EPIC array to analyze ~850,000 CpG probes, helminth-infected individuals had 751 genes with differentially methylated probes (DMPs) compared to uninfected controls (FDR <1%) and of which 72% were hyper-methylated. Gene ontology (GO) analysis identified significant DNA methylation perturbation in the immune system in the IL-4, IFN-γ response, regulation of cytokine production, cell proliferation, mTOR signaling and glycolysis (Figure 1).

**Figure 1:**
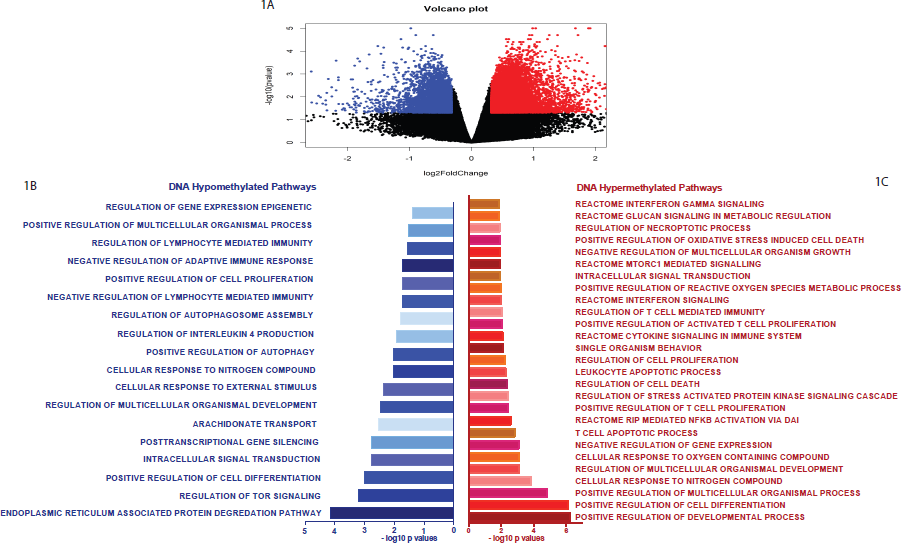
Global CD4^+^ T cell DNA methylation changes during helminth infection. The Illumina DNA methylEPIC array was performed on CD4+ T cells from helminth infected and uninfected individuals. n = 5 per group. A) Volcano plot showing the hypo (blue dots) and hyper (red dots) methylation (β values) in helminth infected individuals compared to uninfected controls. B) Gene ontology pathways enriched in genes with DNA hypo-methylation after helminth infection. C) Gene ontology pathways enriched in genes with DNA hyper-methylation after helminth infection.

To validate these results, targeted DNA methylation of specific genes was evaluated by PCR after methyl-specific endonuclease digestion. The CD4^+^ DNA methylation of transcription factors BATF3, ID2, STAT5A, IRF5, PPARg, RUNX2, IRF4 and NFATC1 were increased by 47%, 48%, 18%, 34%, 52%, 32%, 5% and 60% respectively, above non-helminth infected healthy controls (p< 0.001; Sidak-Bonferroni). Similarly, the CD4^+^ DNA methylation of cytokine or cytokine receptors *IFNGR1*, *TNFS11*, *RELT* (TNF receptor), *IL12RB2* and *IL12B* were increased by 19%, 6%, 12%, 30% and 13% above non-infected controls (p< 0.001; Sidak-Bonferroni; Figure 2a). By contrast, *MAP1LC3*, a gene essential in autophagy, was hypo-methylated 9% in active helminth infection and remained hypo-methylated in remote schistosomiasis (negative by stool and urine O&P and PCR for helminth infection, but serology positive) infection (p< 0.001; Sidak-Bonferroni; Figure 2a)

**Figure 2:**
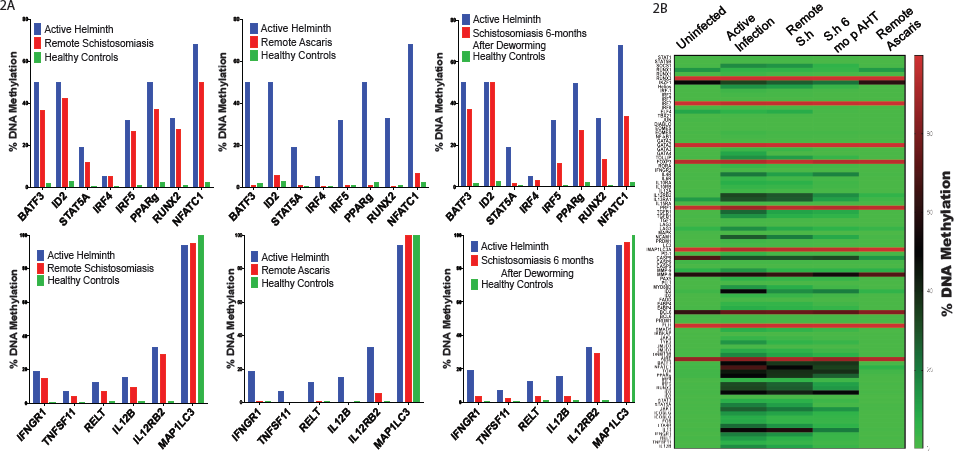
Targeted DNA methylation of transcription factors, cytokines and surface receptors depending on helminth infection. Methyl-specific endonuclease digestion followed by RT-PCR was performed on isolated DNA from CD4^+^ T cells from ascaris-infected, Schistosome-infected infected or uninfected children. Successful deworming was confirmed by negative PCR and microscopy two and six months after anti-helminth therapy. n = 6 per group. A) Bar graph showing percent DNA methylation depending on helminth status. B) Heatmap of the CD4^+^ T cell DNA methylation signature depending on helminth status. (S.h = *Schistosoma haematobium*)

Children with remote *Ascaris* infection had a CD4^+^ DNA methylation signature statistically similar to the healthy control group (Figure 2a). By contrast, children with remote schistosomiasis had CD4^+^ DNA methylation signature statistically similar to the active helminth infection group (Figure 2a). To further evaluate if schistosomiasis induces persistent DNA methylation alterations in CD4^+^ T cells, we repeated the DNA methylation evaluation 6 months after enrollment with negative microscopy and PCR at two and six months to confirm successful deworming. Six months after successful deworming, the DNA methylation of CD4^+^ T cells remained elevated and statistically similar to actively infected individuals (Figure 2a). Comparison between DNA methylation and respective helminth status is shown in the heatmap, Figure 2b. Pathway analysis of these results identified helminth-induced inhibition of the Th1 and IFN-γ signaling pathway (Figure 3 p = 2.97e-33 and p = 6.01e-17, respectively).

**Figure 3:**
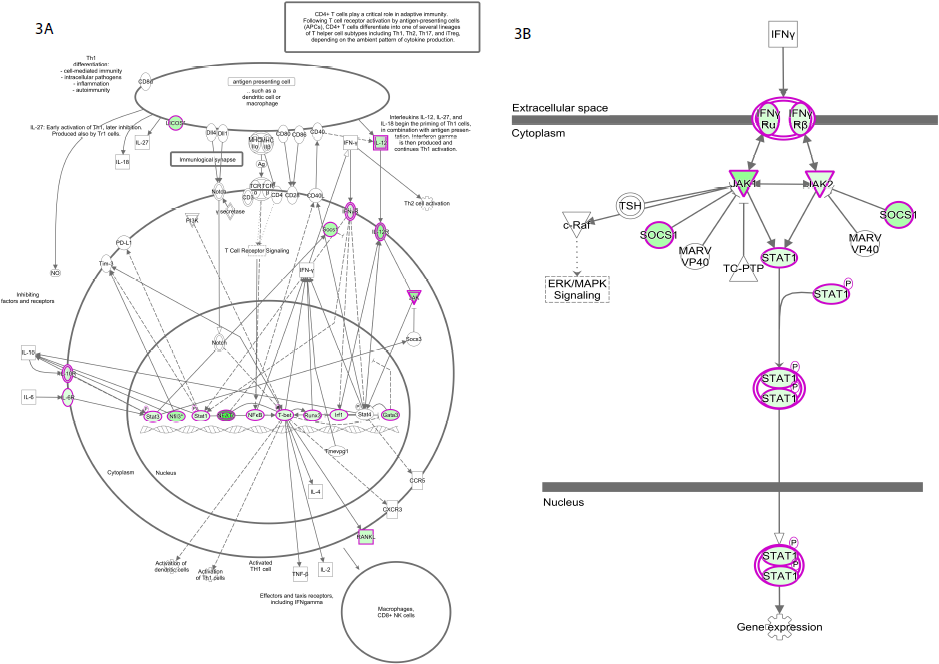
Pathway analysis identified inhibition of the Th1 and IFN-γ signaling pathway by helminth infection. Methyl-specific endonuclease digestion followed by RT-PCR was performed on isolated DNA from CD4^+^ T cells from helminth infected or uninfected children. Genes outlined in pink expressed increased DNA methylation in helminth-infected children compared to uninfected age-matched controls. n = 6 per group. A) Th1 Signaling Pathway. B) IFN-γ signaling pathway.

### Helminth Infection Mutes Up-Regulation of IFN-γ-Inducible

To validate effect of helminth-induced DNA hyper-methylation of the IFN-γ signaling pathway, CD4^+^ T cells were stimulated overnight with 50ng/mL of IFN-γ. The IFN-γ inducible genes specific to CD4^+^ T cells(56) IDO, SERPING1, STAT1, IRF1, FCGR1A/B, GBP1, ICAM1, HLA-DRA, HLA-DRB3 *and HLA-DQA1* were evaluated demonstrating that *IDO* up-regulation was muted by helminth infection (fold up-regulation 12.24 vs 0.02, p = 0.001 Sidak-Bonferroni; Figure 4a). Using a previously described CD4^+^ T cell IFN-γ inducible genes signature(56), helminths induced a 6.7-fold decrease in IFN-γ-stimulated gene up-regulation (*P*<0.05 by Mann-Whitney test; Figure 4b).

**Figure 4:**
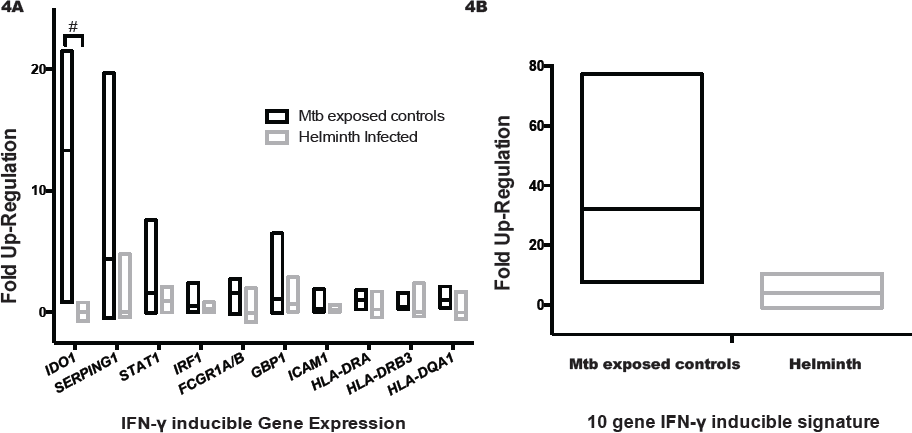
Helminths decrease IFN-γ-inducible gene up-regulation. Peripheral blood mononuclear cells (PBMCs) from recent TB-exposed children were stimulated overnight with IFN-γ (50ng/mL) followed by CD4^+^ T cell selection and RNA isolation. Fold up-regulation was determined by comparing gene expression non-stimulated gene expression to IFN-γ gene expression. n = 8 helminths and 4 controls. A) IFN-γ-inducible gene expression of specific genes. The # symbol denotes Sidak-Bonferroni p< 0.001. B) IFN-γ-inducible gene expression depending on helminth status; P < 0.05, Mann-Whitney. Lines represent median and min to max range.

### Helminth Infection Results in Persistent Immune Phenotypic Perturbations

DNA methylation decreases transcription factor binding affinity, thereby repressing gene expression. To validate the functional impact of helminth infection-induced DNA methylation changes in CD4^+^ T cell, we performed flow cytometry based multi-dimensional immune profiling after *ex vivo* stimulation with TB-specific peptides or BCG sonicate. The characteristic immune alteration induced by helminth infections is an increase in IL-4 production by CD4^+^ T cells, as well as, by other lymphocyte, mast cells, eosinophils and basophils(57, 58). We found that PBMCs from recently-TB-exposed individuals with active helminth infection had an increased frequency of TB-specific (ESAT-6 and CFP-10 stimulation) CD4^+^ T cells producing IL-4 (Wilcoxon signed rank *P* <0.05; Figure 5). The increased frequency of IL-4-producing subsets was associated with a decrease in TB-specific CD4^+^ T cells producing IFN-γ and TNF (Wilcoxon signed rank *p* < 0.05). Thus, DNA methylation alterations in individuals with active helminth infection were associated with a decrease in TB-specific proliferation, as determined by flow cytometry (Wilcoxon signed rank P <0.05). Furthermore, helminth infected individuals had a decrease in the quantity of TB-specific proliferating CD4^+^ T cells (increase in Ki-67 above baseline) producing TNF and IFN-γ (Wilcoxon signed rank P <0.05; Supplemental Figure 1). Since all individuals were BCG vaccinated, the BCG-induced immune response was also evaluated. Notably, individuals with active helminth-infection demonstrated an increased percentage of BCG-induced CD4 T cells producing IL-4 (Wilcoxon signed rank *p*<0.05; Supplemental Figure 2).

**Figure 5:**
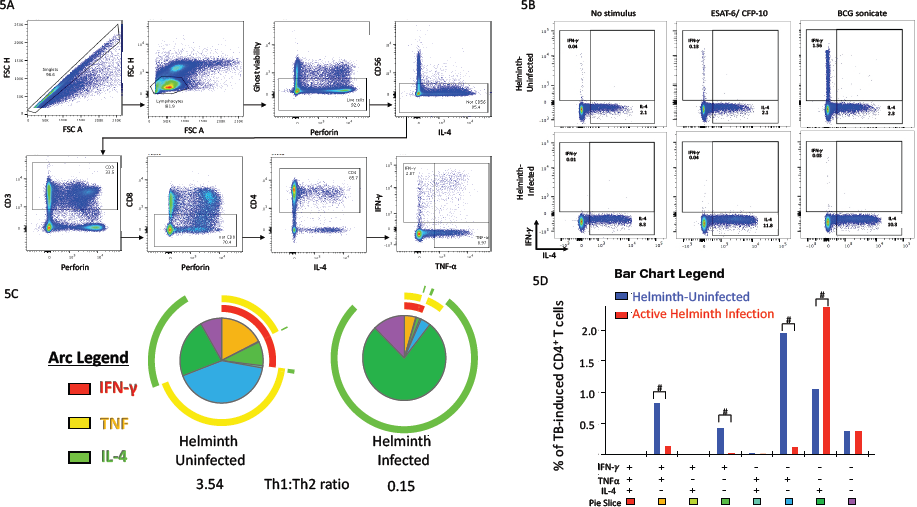
Helminths decrease the *Mycobacterium tuberculosis* (*Mtb*)-specific Th1 CD4+ T cell function with associated increases in Th2. Peripheral blood mononuclear cells (PBMCs) from recent TB-exposed children were stimulated overnight with *Mtb*-specific peptides (ESAT-6 and CFP-10) peptide pools. n = 18 helminth-uninfected and 8 helminth-infected. A) Flow cytometry-gating strategy. B) Representative dot plots of non-stimulated, ESAT-6 and CFP-10 stimulated or BCG stimulated CD4^+^ T cells. C) The overall *Mtb*-specific CD4^+^ T cell response is illustrated as a pie graph with TNF-α (yellow arcs) and IFN-γ (red arcs) and IL-4 (green arcs). Each pie slice corresponds to a discreet functional subpopulation, outlined in D. The Th1: Th2 ratio was determined by the ratio of IFN-γ and TNF-α compared to IL-4. D) The bar graph represents the absolute frequency of CD4^+^ T cells producing, IFN-γ, TNF-α or IL-4. The # symbol denotes Wilcoxon P<0.05 between helminth uninfected and infected groups. Data represent 18 helminth-uninfected and 8 active helminth-infected individuals.

We next evaluated the persistence of helminth-induced immune perturbations on TB-specific and BCG-induced immune responses 6 months after anti-helminth therapy. Successful anti-helminth therapy was documented with negative repeat stool and urine O&P and helminth PCR at 2 and 6 months after study enrollment. For schistosomiasis, but not ascariasis, six months after successful anti-helminthic therapy, the CD4^+^ T cell immune phenotype demonstrated an increase percent of IL-4 and decreased percentage of TNF and IFN-γ antigen-specific producing CD4^+^ T cells compared to uninfected controls (Wilcoxon signed rank *p* <0.05; Figure 6).

**Figure 6:**
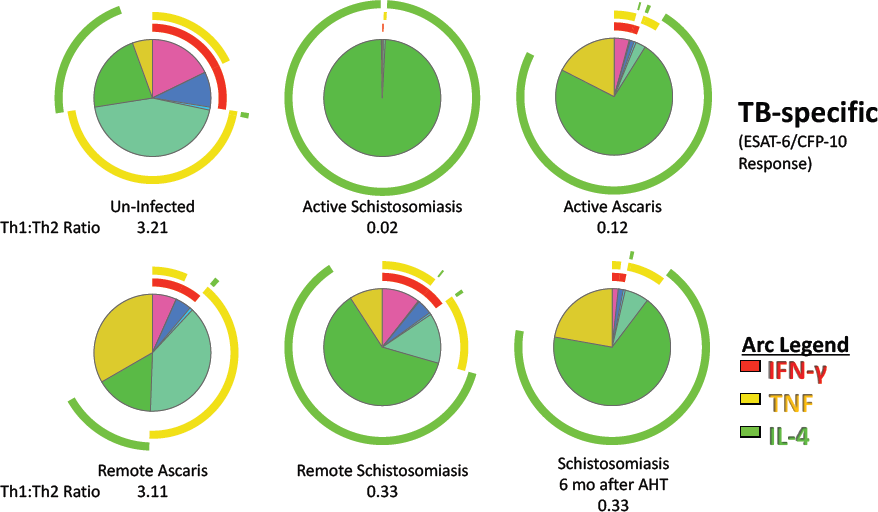
Persistent Th2:Th1 perturbations after schistosomiasis. Children with or without helminth infection had Peripheral blood mononuclear cells (PBMCs) collected at baseline and six-months later. Children with *Ascaris* or *Schistosoma haematobium* were treated with albendazole or praziquantel, respectively. Successful deworming was confirmed by microscopy and PCR at two and six months. The overall *Mtb*-specific CD4^+^ T cell response is illustrated as a pie graph with TNF-α (yellow arcs) and IFN-γ (red arcs) and IL-4 (green arcs). The Th1: Th2 ratio was determined by the ratio of IFN-γ and TNF-α compared to IL-4. Data represent 6 per helminth group and 18 un-infected controls.

Previous reports have established that the supernatant of CD4^+^ T cells incubated with SEA were less effective at forming a mature phago-lysosome with intracellular *Mtb*(32). Consistent with this, monocyte-derived macrophages treated with IL-4 instead of IFN-γ, had a 3.8-fold decreased ability to kill intracellular *Mtb* (Wilcoxon signed rank *p* = 0.039; Figure 7).

**Figure 7:**
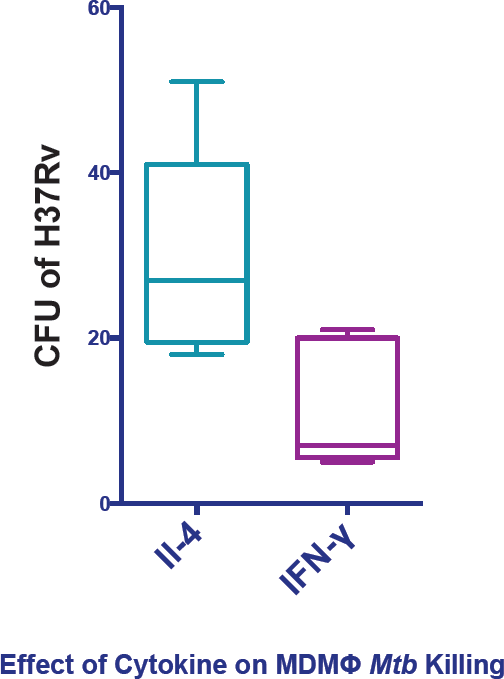
Decreased mycobacterial killing by macrophages after IL-4 polarization. Human monocyte-derived macrophages (MDMΦ) were derived from adherent Peripheral blood mononuclear cells followed by 7 days G-CSF and GM-CSF and 48 hours of polarization with either IL-4 (20ng/mL) or IFN-γ (50ng/mL). In a biosafety level 3 facility, MDMΦs were incubated with H37Rv *Mtb* overnight followed by cell lysis, 1:25 dilution and culture on 7H11J solid culture with colony forming units (CFU) counted at day 21 (*P*<0.05 by Mann-Whitney test). Experiment performed in triplicate from three healthy control donors.

## Discussion

To remain responsive to changing environments, the immune system undergoes epigenetic modifications to refine cell phenotypes(10, 59). CD4^+^ T cells in particular have an epigenetically mediated spectrum of phenotypes that respond to intracellular pathogens (Th1), extracellular parasites (Th2 and Th9), bacteria (Th17 and Th22), and limit autoimmunity (Treg) (59). After encountering a pathogen, both the innate and adaptive immune response retain immunologic memory(2). Here we show that recently TB-exposed, asymptomatic children with helminth infection have CD4^+^ T cell DNA methylation signatures that inhibit the TB-specific and BCG-induced Th1 immune phenotype. Of interest, these CD4^+^ T cell DNA methylation and immune phenotype perturbations persist for at least 6 months in children previously infected with *S. haematobium*, but not in children previously infected with A. *lumbricoides*.

Helminth infection induces a Th2 and immune-tolerant response that subverts cell-mediated immunity to HBV, BCG and cholera(21, 57, 60). The duration of helminth-induced immune changes has been previously evaluated with mixed results. After leaving helminth-endemic areas, refugees experienced decreased eosinophilia, IgE and HLA-DR on CD3^+^ T cells 6-12 months after deworming(60). However other studies have shown that remote *Ascariasis* (as determined by elevated ascaris-specific IgE) is inversely associated with tuberculin skin test (TST) reactivity(61). Further, fourteen months after *in utero* schistosomiasis exposure, children still had a 25-fold decrease in PPD-induced IFN-γ production(21). The mixed results from previous studies are likely due to variability of helminths evaluated and variability of the immune assessments performed. The cohort in the present study consisted of *Ascaris* and *S. haematobium*-infected children who were recently household contacts of individuals with microbiologically confirmed TB. Their helminth status was documented by urine and stool PCR, microscopy and serology. Participants were dewormed and reevaluated 2 and 6 months later to confirm conversion from active to remote helminth status. Similar to the previous studies that evaluated antigen-specific immunity, here the TB-specific and BCG-induced immune response remained shifted away from Th1 immunity six months after successful *S. haematobium* deworming.

There is a paucity of data describing the duration of epigenetic control of immune responses after a host-pathogen interaction. *In vitro*, memory Th1 and Th2 cells retain post-translational epigenetic modifications (H3K4Ac of *IL4* and *IFNG* promoter) for greater than 20 cell divisions(62). In a sepsis model, IL-12 remained epigenetically inhibited for at least 6 weeks(63). *In vivo*, epigenetic alterations persisted 3 months after BCG vaccination(2) and CD8^+^ lymphocytes retain permissive PD1 DNA hypomethylation marks two years after successful antiretroviral therapy(38). Here we show that the schistosomiasis-induced CD4^+^ T cells DNA methylation signature persist at least 6 months after successful deworming for schistosomiasis, but not ascariasis. The continued effect of schistosome infections on the host immune system after treatment is consistent with observations that the pathology associated with schistosomiasis endures beyond infection(64, 65). The difference in the duration of DNA methylation and immune perturbations are likely related to the different life cycles of these organisms. While *Ascaris* is an intraluminal infection and are passed out of the body in tact following treatment, *S.haematobium* worms live in the venous plexus and successful chemotherapy cause a release of adult worm antigens into the bloodstream but the eggs remain imbedded in the tissue causing inflammation that can persist for years after successful treatment(65). For at least some period of time, the eggs continue to induce a Type 2 immune response that is diametrically opposed to the Type 1 immune response directed at intracellular co-infections such as *Mtb*(11, 12, 26, 27, 60). Deworming combined with BCG revaccination was successful at restoring Ag-specific cell-mediated immunity in a previous study of ascaris and hookworm infected individuals(22). This simple strategy should be evaluated in a schistosomiasis-infected population.

After schistosome infection, STAT-6 and JMJD3 mediated alterations in histone H3 lysine-27 (H3K27) tri-methylation induce an alternatively-activated macrophage phenotype(7, 66). Similarly, murine studies have shown that schistosomiasis induces CD4^+^ T cells to shift to a Th2 phenotype associated with DNA hypomethylation of the *IL4* and *Gata3* genes(67). While *IL4* and *Gata3* were not hypo-methylated in this cohort, the gene ontology pathway regulating IL-4 was hypomethylated (Figure 1) with associated increased production of IL-4 (Figure 4). This supports the previous evidence that the Th2 pathway is epigenetically selected and not a default pathway(68).

The importance of the type 1 immune response in controlling mycobacterial infections is highlighted by the rare inherited primary immune deficiencies known as Mendelian Susceptibility to Mycobacterial Disease (MSMDs). MSMDs can be categorized as genetic defects upstream of IFN-γ production (defects in *IL12B, IL12RB, NEMO, ISG15, NEMO*) or defects that block the IFN-γ signaling pathway (*IFNGR1, IFNGR2, STAT1, IRF8*)(30). This work shows that helminths perturb mycobacterial immune response by increasing DNA methylation of both the upstream induction of IFN-γ (*IL12B, IL12RB*) and the downstream IFN-γ signaling pathway (*IFNGR1, IFNGR2, JAK1, STAT1*). DNA hyper-methylation of *IL12B and IL12RB* (Figure 2) in combination with the DNA hypermethylation of Th1 critical transcription factors (Figure 3) are a likely major factor leading to the decreased TB-specific IFN-γ production (Figure 4). Further, by increasing the DNA methylation of the downstream IFN-γ signaling pathway, IFN-γ was less capable of up-regulating the IFN-γ-inducible genes. These experiments focused on the DNA methylation and immune phenotype of CD4^+^ T cells. A critical role of IFN-γ is to induce phenotypic changes in macrophages, specifically to up-regulation of genes necessary for phagolysosome maturation, ROS and NOS, generation of lysosomal acidification. Future studies should evaluate if DNA methylation changes in the IFN-γ signaling pathway block macrophage phagolysosome maturation, ROS and NOS, generation of lysosomal acidification with resultant decreased intracellular *Mtb* killing.

Helminths perturb the immune system and increase the risk of TB progression. Therefore, they are an excellent model to elucidate the immunity necessary to control *M. tuberculosis* infection. In this study, we confirm that *ex vivo* IFN-γ improves *Mtb* killing, but that helminths induce epigenetic DNA methylation changes in CD4^+^ T cells that hamper the IFN-γ production and the downstream intracellular IFN-γ signaling pathway. These findings highlight the importance of persistent DNA methylation perturbations in immune function and provide plausible explanations for increased rate of TB progression in the setting of helminth co-infection. Future studies should evaluate means to reverse helminth-induced detrimental DNA methylation changes to restore mycobacterial immunity. Similarly, longitudinal cohort studies need to be undertaken to evaluate if DNA methylation signatures described here correlate with increased rates of TB progression.

## Disclaimer

The findings and conclusions in this report are those of the authors and do not necessarily represent the views of the Centers for Disease Control and Prevention.

